# China’s Development Assistance for Health on Key Tropical Diseases: A SWOT Analysis

**DOI:** 10.1101/524678

**Authors:** Hong-Mei Li, Wei Ding, Lu-Lu Huang, Xue-Jiao Ma, Ying-Jun Qian, Duo-Quan Wang, Ning Xiao, Ya-Yi Guan, Xiao-Nong Zhou

## Abstract

**Background:** WHO focused and gave priority on ten serious tropical diseases, while China has made remarkable achievements on tropical diseases control. In addition, China has a history of more than 60 years in the health assistance, however, its assistance for tropical diseases started late.

**Methods:** A qualitative questionnaire was distributed to inquiry the opinions of professionals on China’s health assistance on tropical diseases and published articles were searched to collect those data. SWOT analysis, as a tool of qualitative analysis, was used to classify and evaluate the current strengths, the weakness, the opportunities, and the threats of health assistance on tropical diseases in China.

**Results:** Based on SWOT analysis, the internal factors and external environments are obtained. The strengths are focused on China’s achievements on tropical diseases control, surveillance response system of tropical diseases, and human resources of public health; the weakness laid on sustainability of aid projects and funding, applicability of Chinese experience, and lack of composite talents; the opportunities are mainly in the global need of tropical diseases control, China’s health cooperation in Belt & Road and Africa, and the actively participating of international organizations in health assistance; as well as the threats are reflected in the complex international situation, domestic needs of tropical diseases control, and the significant gaps between China and developed countries.

**Conclusion:** The internal strengths and weaknesses of development assistance for health on tropical diseases are clearly presented in the SWOT framework, as well as the external opportunities and threats and corresponding coping strategies. In the era of global health, China should strengthen and improve the health assistance for tropical diseases.

**Author summary:** Tropical diseases are serious infections in tropical and sub-tropical regions, with billions of persons infected and millions of deaths every year, especially in Africa. WHO also called for global efforts to control and eliminate tropical diseases. In the era of global health, development assistance on tropical diseases is important to demonstrate the soft power of national diplomacy, and China has started its health assistance in 1963. In this paper, a qualitative questionnaire and published articles were combined to collect data, and then SWOT analysis was used to analyze the internal factors and external environment, that is the current strengths, the weakness, the opportunities, and the threats of the China’s Health Assistance on key tropical diseases. Based on those results, we put forward the countermeasures and suggestions for the future cooperation of tropical diseases. At the end of this paper, we call on Chinese professionals should make use of their own advantages and actively improve the global tropical diseases control.

## Introduction

China’s development assistance for health began in the early 1950s for Vietnam and North Korea as medical supplies, since then, with the development of international situation and China’s diplomatic strategy, China’s health development mainly experienced three stages [1]. Firstly, the start and foundation (1949-1978), mainly provided free medical supplies and medical aid to African countries; Secondly, adjustment and development period (1979-1999), the main aid measures were to send medical teams, build hospitals, health centers, pharmaceutical factories, health schools and other medical facilities, and carrying out malaria control field in Africa (such as building anti-malaria centers); Thirdly, rapid expansion (2000-to data), in addition to previous measures, the assistance measures were more diversified, including human resources development for health (eg. training for health technicians and officers from developing countries), health related emergency humanitarian assistance and south to south cooperation. It is estimated that China provides about $ 1.5 million annually to Africa [2–3].

In 1963, China dispatched its first medical team to Algeria, which started the sending medical teams to African countries. China’s medical team is the earliest, longest and most distinctive form of assistance in health. Because of the universal and borderless nature, health assistance can stand by the humanitarianism and operate well in the international activities, often achieving the results that cannot achieved by diplomats. Both developed and developing countries attach great importance to health assistance [4–5].

Tropical diseases encompass all diseases that occur solely or principally in the tropics and sub tropics, and often take to refer to infectious diseases that thrive in hot and humid conditions, particularly epidemic in remote areas. WHO focused on ten infectious or parasitic diseases, such as lymphatic filariasis, malaria, schistosomiasis, leishmaniasis, leprosy, tuberculosis, dengue, onchocerciasis, African trypanosomiasis, and chagas disease, as the most common and serious key tropical diseases, which need to be given priority and controlled [6].

China was once a country with severe epidemic of tropical diseases, especially lymphatic filariasis, malaria, schistosomiasis, leishmansis, tuberculosis, leprosy and other diseases that seriously endangered the health of Chinese people. After several generations’ efforts, China has made remarkable achievements in control and elimination of tropical diseases [7]. However, the global situation of tropical diseases control is grim. As the largest developing country, China should actively participate in global cooperation on tropical diseases control and expand its influence in the field of development assistance for health. In this paper, SWOT analysis is performed to analyze the development assistance of health on key tropical diseases in China, so as to provide a base for making relevant planning of development assistance for tropical diseases.

## Materials and Methods

### Data collection

A qualitative questionnaire is distributed to the professionals who participated in the global health training courses on tropical diseases for government officials and managers in 2016, which hold by National Institute of Parasite Diseases, China CDC and supported by UK DFID. The survey consists of five parts, including China’s experience on tropical diseases elimination, the feasibility and challenge of China’s experience spreading to Africa, key works in the next five years and other concerned questions. All participants completed the questionnaire independently, and the staff collected the questionnaire timely. A total of 89 professionals participated in this survey, mainly focused on the terms showed in table 1.

**Table 1.**
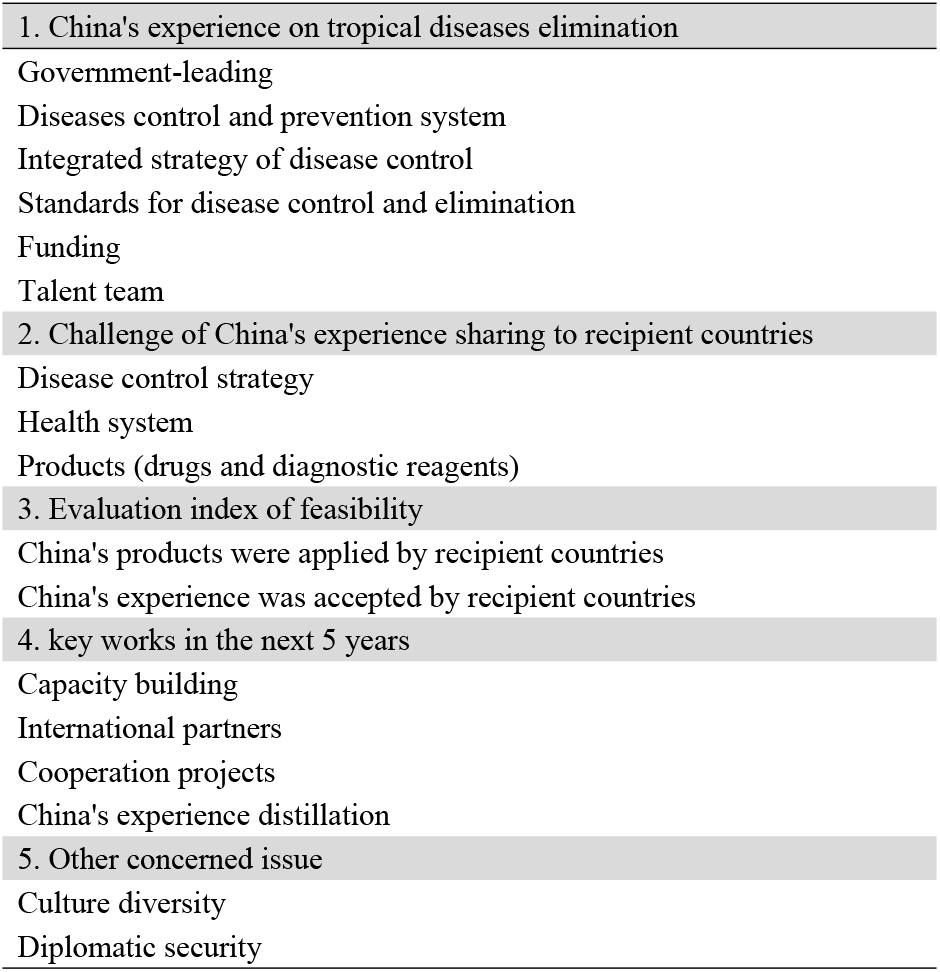
The terms of China’s health assistance for tropical disease focused by professionals in the qualitative questionnaires

### Search strategy

We search articles published on PubMed, CNKI, Google Scholar, and the websites of WHO, UN, CNHC, China CDC, and other related. The search key words were focus on 1) tropical diseases OR any ten key tropical diseases (lymphatic filariasis OR malaria OR schistosomiasis OR leishmaniasis OR leprosy OR tuberculosis OR dengue OR onchocerciasis OR African trypanosomiasis OR chagas disease); 2) health assistance OR health aid; 3) global health; 4) the terms focused by Chinese professionals shown in table 1. Literatures both in English and Chinese are enrolled, and were identified by manual screening.

### Qualitative analysis

All questionnaires were read, coded, extracted, and categorized into different classifications. Combined with the results of questionnaires and articles, a SWOT analysis was used for qualitative analysis to evaluate the current performance of health assistance on tropical diseases to develop strategic plans for Chinese government improving future operations, which included four components, the current strengths, the weakness, the opportunities, and the threats. Those first two are also classified as internal factors and the latter two are external environments.

## Result

### Strengths of health assistance in key tropical diseases Achievements of key tropical diseases control in China

Over the past decades, thanks to the attention paid by the Chinese government and the application of comprehensive strategies on tropical diseases control, China has achieved unprecedented development and achievements in the control and elimination of tropical diseases (Table 2). In 2007, China successfully eliminated lymphatic filariasis, and was recognized by WHO, becoming the first country to eliminate lymphatic filariasis in 83 endemic countries and regions in the world [8]. This is another achievement in the field of public health since China announced eradicating smallpox and polio. China has effectively controlled the epidemic of malaria, and the epidemic intensity and extent of malaria have decreased significantly. The transmission of falciparum malaria is successfully blocked in the central of China, and 99.9% of the report cases are imported malaria cases [9]. In 2016, China reached the standard of schistosomiasis transmission control, which greatly accelerated the progress of schistosomiasis control in China [10–11]. In 1958, most of the endemic areas in China have basically eliminated visceral lershmaniasis, which is now mainly prevalling in northwestern China [7, 12]. Leprosy is generally at a low epidemic level in China, and the key endemic areas are in southwestern provinces [13]. For the tuberculosis prevention, although the epidemic situation in China is still severe, some achievements have been made [14–15]. In addition, although China is not the epidemic area of dengue, African trypanosomiasis, onchocerciasis and chagas disease, imported cases are often reported, especially dengue [16], which should be paid more attention. To further consolidate the achievements of tropical diseases control, China has put forward the goal of eliminating malaria nationwide by 2020 and eliminating schistosomiasis by 2025.

**Table 2.** Prevalence of ten key tropical diseases in China and the world

| Name | Pathogen | Distribution and prevalence in China | Global distribution and Prevalence |
| --- | --- | --- | --- |
| lymphatic filariasis | *Wuchereria bancrofti*,<br>*Brugia malayi*, *Brugia timori* (parasite) | Lymphatic filariasis has been eliminated in 2007 [8]. | According to WHO, 856 million people in 52 countries worldwide threatened by lymphatic filariasis. In 2000 over 120 million people were infected, with about 40 million disfigured and incapacitated by the disease [28]. |
| malaria | *Plasmodium* (parasite) | In 2016, a total of 3321 cases of malaria were reported, including 3 cases of local infection and the rest were imported cases. It is mainly distributed in yunnan, sichuan, jiangsu, guangxi and shandong [9]. | In 2017, there were an estimated 219 million cases of malaria in 90 countries, and Africa was home to 92% of malaria cases and 93% of malaria deaths [29]. |
| schistosomiasis | *Schistosoma* (parasite) | In 2016, it is estimated 54454 cases infected schistosomiasis, and China was generally in a low prevalence state [10-11]. | Schistosomiasis transmission has been reported from 78 countries. Estimates show that at least 206.4 million people required preventive treatment for schistosomiasis in 2016, and at least 91.4% of those requiring treatment for schistosomiasis live in Africa [30]. |
| leishmaniasis | *Leishmania* (parasite) | Visceral leishmaniasis mainly epidemic in xinjiang, gansu and sichuan provinces, where the reported cases accounts for more than 97% of the total cases of China [7, 12]. | An estimated 700 000 to 1 million new cases of leishmaniasis and 20 000 to 30 000 deaths occur annually [31]. |
| leprosy | *Mycobacterium leprae* (bacillus) | In 2015, there were 3230 cases of leprosy in China, and the prevalence rate was 0.235/100000, and the provinces with high | It is reported 173 358 cases of leprosy at the end of 2016, prevalence rate corresponds to 0.29/10,000. Elimination of leprosy as public health problem (defined as a registered |
|  |  | prevalence were yunnan, guizhou, sichuan and guangdong [13]. | prevalence of less than 1 case per 10 000 population) was achieved globally in 2000 [32]. |
| tuberculosis | *Mycobacterium tuberculosis* (bacteria) | In 2010, WHO estimated the annual cases of tuberculosis in China was 1 million, mainly in guangdong, henan, hubei, sichuan, chongqing, shaanxi, jiangxi and xinjiang [14-15]. | Tuberculosis (TB) is one of the top 10 causes of death worldwide. In 2017, 10 million people fell ill with TB, and 1.6 million died from the disease. In 2017, the largest number of new TB cases occurred in the South-East Asia and Western Pacific regions, with 62% of new TB cases, followed by the African region, with 25% of new TB cases. In 2017, 87% of new TB cases occurred in the 30 high TB burden countries. Eight countries accounted for two thirds of the new TB cases: India, China, Indonesia, the Philippines, Pakistan, Nigeria, Bangladesh and South Africa [33]. |
| dengue | *Dengue virus* (virus) | No local dengue epidemic, however, imported cases have appeared in guangdong, yunnan, fujian, zhejiang, guangxi and taiwan provinces [16]. | It is estimated that 390 million dengue infections per year, of which 96 million manifest clinically. Now endemic in more than 100 countries, and the America, South-East Asia and Western Pacific regions are the most seriously affected [34]. |
| onchocerciasis | *Onchocerca volvulus* (parasite) | Non-endemic areas of onchocerciasis | More than 99% of onchocerciasis infected people live in 31 African countries. The disease also exists in some foci in Latin America and Yemen. It is estimated 20.9 million *Onchocerca volvulus* infections worldwide. More than 99% infected people live in Africa [35]. |
| trypanosomiasis | *Trypanosoma brucei gambiense*, *Trypanosoma brucei rhodesiense*<br>(parasite) | Non-endemic areas of trypanosomiasis | Trypanosomiasis occurs in 36 sub-Saharan Africa countries. In 2009 the number reported dropped below 10 000 for the first time in 50 years, and in 2015 there were 2804 cases recorded [36]. |
| chages disease | *Trypanosoma cruzi*<br>(parasite) | Non-endemic areas of chages disease | Chagas disease is found mainly in endemic areas of 21 Latin American countries. About 6 million to 7 million people worldwide, mostly in Latin America [37]. |

### Surveillance and response system on tropical diseases

Since 2004, China launched and built the information network for surveillance and monitoring of legal infectious diseases, using the internet technology to build the national information platform of monitoring infectious cases, with the functions of dynamic monitoring, real-time reports and statistics, which links the health administrative departments, health center and related organization from four levels (county level – province level – county level – township level) [17–19]. At present, this information system has covered all county level administrative divisions, and already became the largest surveillance system of infectious diseases in the world [20]. A total of 39 species of infectious diseases have been covered in this system, including lymphatic filariasis, malaria, schistosomiasis, tuberculosis, leprosy, leishmaniasis, dengue, that is to say, in addition to the 3 tropical diseases non-endemic in China (trypanosomiasis, onchocerciasis and chagas disease), the other 7 key tropical diseases have been covered in this surveillance system. Surveillance and response system can monitor outbreaks dynamically and implement the case management timely, not only improving the accuracy of disease surveillance, but also enhancing the capability of early detection, which pays a crucial role in the progress of diseases prevention, control and elimination. In addition, China has also established the information system for case management of parasite diseases [21–22], including the important parasite diseases epidemic in China, such as malaria, schistosomiasis, filariasis and so on.

### Human resources of public health

At present, the medical university in China usually have the disciplines of public health and preventive medicine, and set specialized, undergraduate and postgraduate majors. Among them, many schools offer the major of parasitology or take parasitology as the research direction. Researchers from national, provincial, municipal and county centers for diseases control and prevention as well as the specialized institutes of parasite diseases have accumulated rich practical experience through years participating in different types of prevention and control projects on tropical diseases, which had improved the capacity of tropical diseases control greatly. Especially in recent years, with the upsurge of global health, many famous universities have set up the new global health school, which will injects new vitality into the cultivation of international health talents. The establishment of the Chinese Consortium of University for Global Health (CCUGH), the China Global Health Network (CGHN) and Chinese Society of Global Health (CSGH) also contribute to the building of the global health think tank and leading the talent cultivation. In addition, the China-UK global health project has put the concept of China’s health development assistance into practice, and also trained a number of global health managers and practitioners for China.

### Weaknesses analysis of health assistance in key tropical diseases Sustainability of the projects and funding of health assistance

The health assistance on tropical diseases in China is mainly including carrying out cooperation projects and personnel exchanges from both sides, and those activities are mostly dependent on external funding, such as the support from WHO, the global fund, the gates foundation, and the government funds of developed countries (United Kingdom and Australia), and so on. However, if those funding is used up, the projects is end, then the China’s assistance is in a dilemma, or even stagnation. So the sustainability of China’s assistance in health financing and the continuity of aid projects on tropical diseases need to be strengthened. In addition, China has not established a unified management mechanism of health development assistance for tropical diseases, and the cumbersome customs formalities for medical equipment and medicines, as well as the improper transport and use of drugs, have also affected the assistance process to some extent. China’s assistance model for tropical diseases is relatively simple and the assistance force is relatively dispersed, making it difficult to form a synergy, which are leading to the impact on health system of the recipient countries is limited.

### The feasibility of China’s experience

The aid priority of OECD (Organization for Economic Co-operation and Development) countries focused on the specific diseases, in contrast, China’s aid focus on constructing health facilities, dispatching medical teams, and training medical workers, and very few programs for specific diseases. In general, China’s development assistance for health is given priority to “hard” health facilities, however, the investment is inadequate to the “soft” health knowledge, experience and product, such as supplying drugs, devices and vaccine, construing diseases diagnosis standards, and professional health advising [20]. It is claimed that due to lack of systematic data to assess China’s aid, it is difficult to value those projects. So the effect and reliability of China aid are difficult to define, it is still lack the data for feasibility and applicability of China’s experience [23].

### Lack of integrated talent

Along with the progress of globalization speeds up, the traditional training model of public health and preventive medicine could not meet the demand of the development assistance for health, needs to be combined with global health. However, the discipline of global health in its infancy, there is no mature model in the world. The traditional public health is only built on the health sector alone to solve medical problems, just from the medicine itself to find and solve problems, there is a significant differences with global health. Professionals of global health need to master not only medicine, law, economics, management science and informatics and other multi-disciplinary knowledge, but also need to understand culture, religion and health systems of different countries [24]. More importantly, those professionals could owned the ability to discover, analyze and solve cross-border health problems with globalization perspective and solve the health globalization problems using inter-disciplinary and multi-sector resource.

### Opportunities of health assistance in key tropical diseases The demand of global tropical diseases control

In 2015, the UN sustainable development goal proposed to eliminate epidemics such as AIDS, tuberculosis, malaria and neglected tropical diseases by 2030 [25], covering those 10 key tropical diseases, among which 8 are also neglected tropical diseases. As early as 2012, WHO issued the global strategy for the prevention and control of neglected tropical diseases, and signed the London declaration, proposing the goal of eliminating or controlling neglected tropical diseases by 2020 [26–27]. Thus it can be seen that tropical diseases are seriously harmful and widespread, and global action is urgently needed. According to the WHO report (table 2), lymphatic filariasis is endemic in 52 countries with over 856 million people threatened [28]. Malaria is endemic in 90 countries. In 2017, there were 219 million cases of malaria worldwide, and 92% of them were from Africa, which accounted for 93% of the total malaria deaths [29]. Schistosomiasis is endemic in 78 countries, with 206.5 million people infected worldwide, and at least 92% of those requiring treatment living in Africa [30]. There are about 0.7-1 million new cases of leishmaniasis worldwide each year, and about 20-30 thousand deaths [31]. In 2000, the goal of elimination leprosy as a public health problem was achieved globally (defined as a prevalence lower than 1 case per 10 thousand people) [32]. Tuberculosis is one of the top 10 causes of death worldwide. In 2017, about 10 million people worldwide fell ill with tuberculosis, and 87% of new TB cases occurred in the 30 high TB burden countries [33]. About half of the world’s population is at risk of dengue, mostly in tropical and subtropical areas [34]. More than 99% of onchocerciasis cases occur in 31 African countries [35]. African trypanosomiasis is endemic in 36 south Sahara African countries [36]. The number of chagas infected is estimated to be between 6 million to 7 million, mainly in 21 Latin American countries [37].

### “Belt & Road” initiative and China-Africa health cooperation

In recent years, China’s assistance to African countries has been increasing, and China has become one of the top 10 global health donors in Africa [38]. Chinese government has strengthened its health assistance work with Belt & Road countries and African countries through a series of measures. In 2013, China began to put forward the Belt and Road initiative, promoting “policy communication”, “facility connectivity”, “unimpeded trade”, “financing integration” and “people-to-people bonds” as five major goals. Health cooperation is the key to opening up “people-to-people bonds” [39]. In October 2016, China formulated the plan for “healthy China 2030”, which comprehensively promote international cooperation of health. In January 2017, China and WHO signed the “Belt & Road” MoU on health cooperation, and the implementation plan in July 2017, including the control of tuberculosis, malaria, and schistosomiasis. In August 2018, Beijing Action Plan was announced in the Forum on China-Africa Cooperation, in which, infectious diseases such as tuberculosis, malaria and schistosomiasis were included in the key diseases of China to support African countries, and committing to support the establishment of the African CDC [40].

### The development of biotechnology and medicine bringing new diagnostic and therapeutic tools

The application of new technologies represented by biotechnology, such as pathogenic biology, genomics, proteomics and bioinformatics, has effectively revealed the biological characteristics of tropical diseases and promoted the pathogenesis research, new prevention technologies and drug development. Those further promoted the development of tropical diseases prevention and control. For example, the development of safe and effective vaccines is the key requirement for global malaria control, while biotechnology has effectively promoted the development process of vaccine [41–42]. Rapid detection reagent strips, ELISA and PCR have been successfully applied to malaria diagnosis. On the other hand, the development of new drugs has greatly promoted the process of global tropical disease prevention and treatment, such as: artemisinin combining with other drugs is the effective ways to treat malaria. In addition, international pharmaceutical companies are also actively involved in the global tropical diseases control.

### International organizations keened on development assistance for health

On the stage of global development assistance for health, there are lots of active international organizations [43]. In recent years, the cooperation between China and WHO has gradually increased, and the WHO collaborating center for tropical diseases has been established in 2016, which had laid an important foundation for strengthening cooperation with WHO in tropical diseases. In addition, bilateral agencies and international foundations are also keen on tropical diseases assistance [44]. UK DFID mainly supports the infectious diseases control (including malaria and schistosomiasis), maternal and child health, community health services and capacity building; the Bill & Melinda Gates Foundation is committed to prevent infectious diseases including polio, malaria, AIDS, diarrhea, pneumonia and tuberculosis; Global Fund are focus on prevent and treat AIDS, tuberculosis and malaria. Public health problems such as tropical diseases have crossed national borders, joint action and cooperation among organizations are needed to deal with the challenges of health globalization in carrying out health development assistance.

### Threats of health assistance in key tropical diseases The complex international situation and needs of domestic disease control

With the agglomeration of globalization, unceasing changes on the international economic structure and restructuring of global governance structure, health assistance has gradually become an important way for countries to carry out health diplomacy. On the one hand, the international situation is increasingly complicated, developed countries are the old-fashioned aid countries, and the strength of the emerging economies continues to grow, making the competition among the donor countries fierce [1]. On the other hand, some domestic parties and armed conflicts in the recipient countries have led to political and economic instability, making health assistance more focused on diplomatic security and exacerbating the complexity of aid [45]. China not only has to face the complicated international situation, but also address domestic health needs, especially the problems of insufficient health resources and serious diseases burden in the central and western regions, for example, leishmaniasis in Xinjiang, Qinghai and Sichuan is very serious.

### Significant gaps between China and foreign countries

First of all, developed countries have a long history of experience in providing health assistance. In particular, since 2006, seven OECD countries such as Switzerland, the United Kingdom, the United States, Japan, Norway, France, and Germany have successively issued national global health strategies [46], by which the internal unity of health and foreign policy has been achieved, so as to the health and foreign affairs departments have been able to defend national interests on the international stage. However, China has not yet introduced a national global health strategy and lacks top-level design and planning. As a representative of emerging economies, China will play an important role in South-South health cooperation, South-North health cooperation, Belt & Road health cooperation and global health governance. Therefore, China also needs to formulate a national health strategy as soon as possible to better guide various actors in their commitment to people’s health and global health cooperation. In addition, China’s health assistance concepts and models, related technologies and products exported, and health diplomacy capabilities are more or less challenged by international rules formulated by developed countries, and there is an urgent need to improve the cooperation level and capacity of China’s health assistance.

The results of SWOT analysis were obtained in picture 1.

**Picture 1.**
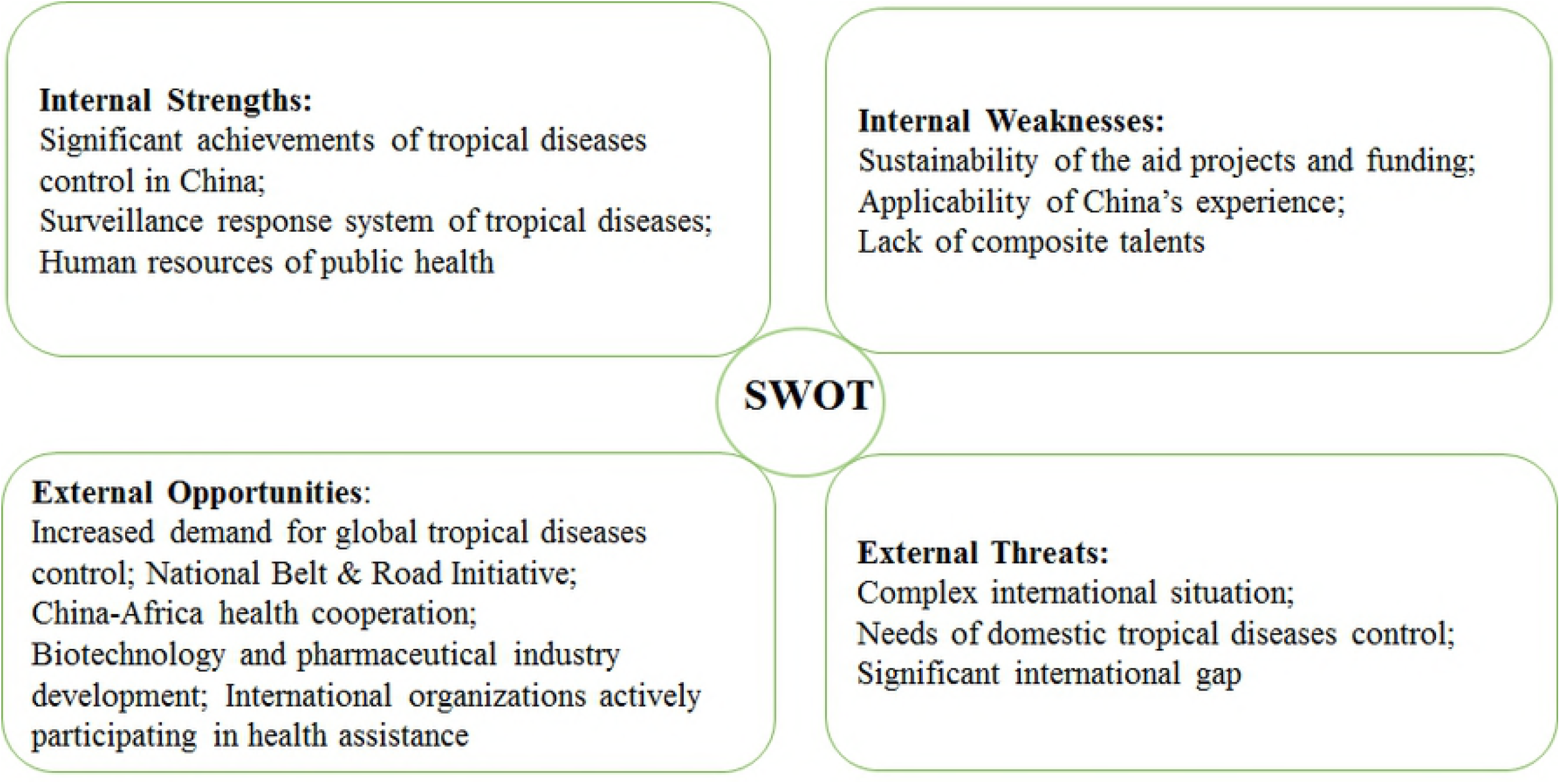
SWOT analysis of China’s development assistance for health on tropical diseases

## Discussion

SWOT analysis is a useful tool for understanding the current strengths and weaknesses, and for identifying the opportunities and threats. In this paper, the results of SWOT analysis not only provided a framework for evaluating the benefits and limitations of current situation, but also laid a base to bring forward four strategic choices for China’s development assistance for health on tropical diseases.

### Firstly, SO (Strengths-Opportunity) Strategy: Seize the opportunity and develop its advantages

China has accumulated rich experience in the tropical diseases control. It is necessary to fully summarize China’s own experience, eg: how to carry out epidemiological investigations, how to find out the prevailing situation, conduct vector monitoring, conduct surveys on intermediate host, and carry out disease monitoring. The experience of monitoring, interventions, diagnosis and treatment will be refined, and those will be integrated into a set of technologies that can be directly applied. The Chinese story will be well explained. We should make full use of these experiences and wisdom to actively contribute to the Chinese discourse and strength. The disease surveillance response system is a key point on tropical diseases control in China. Rational use of the surveillance response system can timely grasp the patient’s personal information, the pathogens, place of residence and the infection situation, which is conducive to timely control the epidemic. It is necessary to widely publicize and promote the connotation and role of the surveillance response system, and emphasize the importance of the surveillance response system for tropical diseases control. At the same time, it is also necessary to make full use of biotechnology to develop new tools and methods for tropical diseases detection and therapeutic drugs. In recent years, drug resistances have been found in many areas, especially malaria. To restrain the rising malaria drug resistance situation, new antimalarial drugs must be developed, which is also crucial for diseases control.

### Secondly, ST (Strengths-Threats) Strategy: Build on Strengths and resolve threats

Peaceful development and international cooperation have become the main theme of the times. China should actively participate in global health governance and demonstrate its image as a responsible country. Tropical diseases have crossed the traditional geographical and national boundaries. In response to its prevalence and outbreaks, it is difficult for a country to be independent, and only through cooperation can benefit. On the contrary, it will be threatened by the outbreak of epidemic and endanger the people health. As the largest developing country, China should make use of its advantages to develop assistance with other developing countries (Southeast Asian and African countries) while doing its own disease control. First of all, China should formulate its own global health strategy, joint the health, diplomacy, education and other related departments, to carry out tropical diseases assistance together. China also need to continuously innovate the assistance models, improve aid management systems and working mechanisms, and enhance aid technologies and level, those can contribute to carry out health assistance and cooperation projects of tropical diseases better. For example, helping recipient countries to set up tropical diseases control plans, export products such as drugs and diagnosis reagents, and establish standardized aid evaluation indicators.

### Thirdly, WO (Weaknesses-Opportunities) Strategy: Use opportunities to change disadvantages

China has repeatedly mentioned the aid commitments to Africa. Both the Belt & Road initiative and China-Africa health cooperation contained the tropical diseases assistance. In addition, international organizations and foundations are seeking the partners, expecting carrying out joint efforts on health assistance, therefore, China’s experience of tropical diseases control is highly compatible with the global needs of tropical diseases prevention [51], but it is unavoidable that China’s experience directly applies to African countries are challenged due to the differences in culture, language, religion, policy system and disease characteristics. How to take advantage of opportunities to deal with challenges. Firstly, we should strive the national and international support, launch a number of cooperation projects with national organizations and international foundations and establish pilots with innovative diseases control models in recipient countries, through which we can explore how to share Chinese experience and technologies to local conditions. Secondly, we need to cultivate global comprehensive talents for health. In the era of global health, requirements for the professional of tropical diseases control are not only equipped rich public health knowledge, but also possessed a strong international diplomatic ability. The global expert team of health aid and tropical diseases should be strengthened through the introduction, training and exchanges of talents.

### Fourthly, WT (Weaknesses-Threats) Strategy: Draw lessons from experience to make up for the shortfall

The emergence of global health has accelerated the new trend in global health governance. Faced with the complicated international situation, China should strengthen the management of tropical disease assistance projects. Before the project is launched, China should fully carry out investigates and survey. After the project is launched, China should emphasize dynamic supervision, coordinate and solve problems timely, and implement progress management and effect evaluation. Developed countries have relatively mature health assistance systems. For example, UA AID mainly provides health assistance in health policy formulation, high-level political cooperation, technical support and personnel training. China should learn from the advanced international health assistance methods and practices, and use those successful experience to guide the future development. However, development and risk coexist. The needs of global tropical diseases control develop a space for the development of China’s health assistance, similarly, risks are also included. For example, instability of funding sources for aid projects that may be brought about by the freeze of international funds, as well as the tensions in recipient countries may jeopardize the safety of aid workers. Those requires keeping up with the international situation, predicting ahead, and taking preventive measures.

Through the above analysis, the strategic choices for the development assistance for health on tropical diseases can be constructed, as shown in Table 3.

**Table 3.** Four strategic choices of the development assistance for health on tropical diseases in China based on SWOT analysis

| Strategic Combination | Concrete Measures |
| --- | --- |
| **SO Strategy** | Take advantage of the surveillance response system;<br>Actively share Chinese experience;<br>Develop new technologies and drugs for tropical diseases |
| **WO Strategy** | Strive for domestic and foreign aid projects and funds;<br>Share Chinese experience according to local conditions;<br>Cultivate comprehensive talents |
| **ST Strategy** | Development of China's global health strategy timely;<br>Innovative aid model;<br>Improved the aid levels |
| **WT Strategy** | Strengthen project management;<br>Learn from the international leading health assistance experience;<br>Predict the risks of tropical diseases aid projects |

## Conclusion

Based on the SWOT analysis, the internal strengths and weaknesses of development assistance for health on tropical diseases can be clearly seen in the era of global health, as well as the external opportunities and threats and corresponding coping strategies. In order to further improve the health assistance for tropical diseases, we need to concentrate on refining the set of Chinese technologies and experience for tropical diseases, formulating China’s health assistance strategy, conducting cooperation pilot research projects in recipient countries, and strengthening the construction of global health assistance teams.

In conclusion, if a country is not involved in the prediction, prevention and improvement of global health problems, it is not only the health of the country’s population that is threatened, but also the national economy and security. The arrival of the global health era has brought new opportunities for the field of health development assistance. Tropical disease professionals should make use of their own advantages, rely on domestic and international health cooperation trends, improve their capacities, innovate aid models and management methods, and actively promote the progress of global tropical diseases control and realize the value of Chinese experience.

## Competing interests

The authors have declared that no competing interests exist.

## Notes

* Supported by China-UK Global Health Support Programme funded by UK DFID (No. GHSP-CS-0P2-02). The funder had no role in study design, data collection and analysis, decision to publish, or preparation of the manuscript.

